# GWAS in 446,118 European adults identifies 78 genetic loci for self-reported habitual sleep duration supported by accelerometer-derived estimates

**DOI:** 10.1101/274977

**Authors:** Hassan S Dashti, Samuel E Jones, Andrew R Wood, Jacqueline M Lane, Vincent T. van Hees, Heming Wang, Jessica A Rhodes, Yanwei Song, Krunal Patel, Simon G Anderson, Robin Beaumont, David A Bechtold, Jack Bowden, Brian E Cade, Marta Garaulet, Simon D Kyle, Max A Little, Andrew S Loudon, Annemarie I Luik, Frank AJL Scheer, Kai Spiegelhalder, Jessica Tyrrell, Daniel J Gottlieb, Henning Tiemeier, David W Ray, Shaun M Purcell, Timothy M Frayling, Susan Redline, Deborah A Lawlor, Martin K Rutter, Michael N Weedon, Richa Saxena

## Abstract

Sleep is an essential homeostatically-regulated state of decreased activity and alertness conserved across animal species, and both short and long sleep duration associate with chronic disease and all-cause mortality^1,2^. Defining genetic contributions to sleep duration could point to regulatory mechanisms and clarify causal disease relationships. Through genome-wide association analyses in 446,118 participants of European ancestry from the UK Biobank, we discover 78 loci for self-reported sleep duration that further impact accelerometer-derived measures of sleep duration, daytime inactivity duration, sleep efficiency and number of sleep bouts in a subgroup (*n*=85,499) with up to 7-day accelerometry. Associations are enriched for genes expressed in several brain regions, and for pathways including striatum and subpallium development, mechanosensory response, dopamine binding, synaptic neurotransmission, catecholamine production, synaptic plasticity, and unsaturated fatty acid metabolism. Genetic correlation analysis indicates shared biological links between sleep duration and psychiatric, cognitive, anthropometric and metabolic traits and Mendelian randomization highlights a causal link of longer sleep with schizophrenia.

---

Research in model organisms (reviewed in ^3,4^) has delineated aspects of the neural-circuitry of sleep-wake regulation^5^ and molecular components including specific neurotransmitter and neuropeptide systems, intracellular signaling molecules, ion channels, circadian clock genes and metabolic and immune factors^4^, but their specific roles and relevance to human sleep regulation are largely unknown. Prospective epidemiologic studies suggest that both short (<6,7 hours per night) and long (>8,9 hours per night) habitual self-reported sleep duration associate with cognitive and psychiatric, metabolic, cardiovascular, and immunological dysfunction as well as all-cause mortality compared to sleeping 7-8 hours per night^6–12^. Furthermore, chronic sleep deprivation in modern society may lead to increased errors and accidents^13^. Yet, whether short or long habitual sleep duration causally contributes to disease initiation or progression remains to be established.

Habitual self-reported sleep duration is a complex trait with an established genetic component (twin- and family-based heritability (*h*^2^) estimates =9-45%^14–20^). Candidate gene sequencing in pedigrees and functional validation of rare, missense variants established *BHLHE41* (previously *DEC2*), a repressor of *CLOCK*/*ARNTL* activity, as a causal gene ^21,22^, supporting the role of the circadian clock in sleep regulation. Previous genome-wide association studies (GWAS) in up to 128,286 individuals identified association of common variants at or near the *PAX8* and *VRK2* genes^20,23,24^.

Here, we extend GWAS of self-reported sleep duration in UK Biobank to discover 78 loci, test for consistency of effects in independent studies of adults and children/adolescents, determine their impact on accelerometer-derived estimates, perform pathway and tissue enrichment to highlight relevant biological processes and explore causal relationships with disease traits.

Among UK Biobank participants of European ancestry (*n* =446,118), mean self-reported habitual sleep duration was 7.2 hours per day (**Supplementary Table 1**). GWAS using 14,661,600 imputed genetic variants identified 78 loci for self-reported habitual sleep duration (*P* <5x10^-8^; **Figure 1a, Supplementary Table 2, Supplementary Figure 1a, Appendix 1**). Individual signals exert an average effect of 1.04 [95% CI: 1.00-1.07] minutes per allele, with the largest effect at the *PAX8* locus, with an estimate of 2.44 [2.14-2.75] min per allele, consistent with previous reports^20,23,24^. The 5% of participants carrying the most of the 78 sleep duration-increasing alleles self-reported 22.2 [22.1-22.3] minutes longer sleep duration compared to the 5% carrying the fewest. The 78 loci explain 0.69% of the variance in sleep duration, and genome-wide SNP-based heritability^25^ was estimated at 9.8 (0.001)%.

**Figure 1a.**
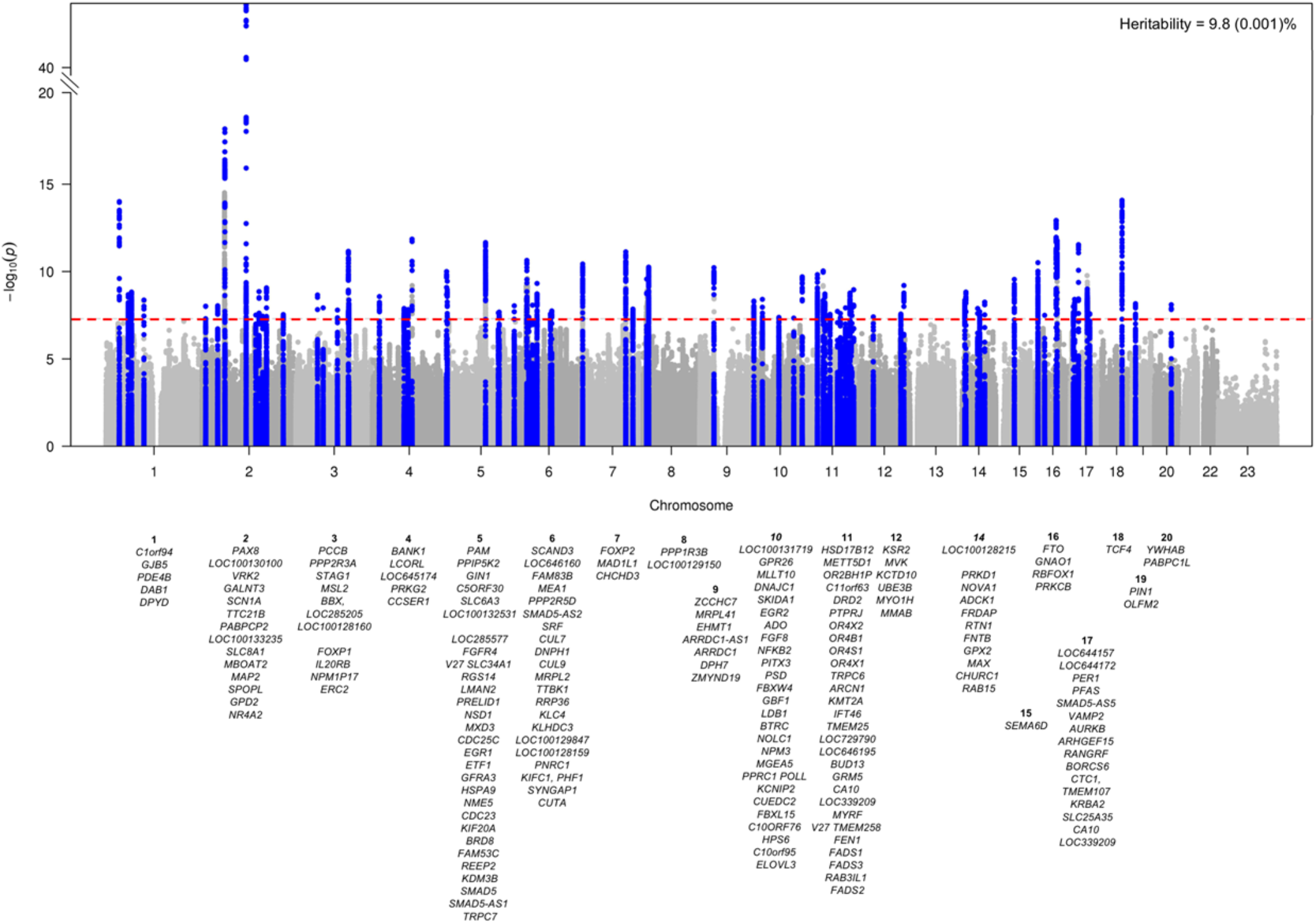
Plots for genome-wide association analysis results for A) sleep duration and B) short and long sleep. Manhattan (a) and Miami (b) plots show the -log_10_ *P* values (y-axis) for all genotyped and imputed SNPs passing quality control in each GWAS, plotted by chromosome (x-axis). Horizontal red line denotes genome-wide significance (*P* =5 x 10^-8^). **A.** Sleep duration (*n* =446,118)

Sensitivity analyses indicated that the 78 genetic associations were largely independent of known risk factors (**Supplementary Table 3**). Effect estimates at 18/78 loci were attenuated by 15-30% upon adjustment for frequent insomnia symptoms, perhaps reflecting contribution to an insomnia sub-phenotype with physiological hyperarousal and objective short sleep duration^26^ (**Supplementary Table 3**). Interestingly, no further signal attenuation was observed when accounting for BMI at rs9940646 at *FTO*, the established BMI-associated signal (*r^2^* =0.81 with rs9939609^27^ and where the higher BMI allele associates with shorter sleep duration). Conditional analyses identified 4 secondary association signals at the *VRK2*, *DAT1 (SLC6A3)*, *DRD2*, and *MAPT* loci (**Supplementary Table 4**). Effect estimates were largely consistent in GWAS excluding shift workers and those with prevalent chronic and psychiatric disorders (excluding *n* =119,894 participants) (**Supplementary Table 2,5, Supplementary Figure 1b,2, Appendix 1**). GWAS results were similar for men and women [*r*_g_ (SE) =0.989 (0.042); *P*<0.001] (**Supplementary Tables 6, Supplementary Figure 1c,1d,3**).

Separate GWAS for short (<7 hours; *n* =106,192 cases) and long (≥9 hours; *n* =34,184 cases) sleep relative to 7-8 hours sleep duration (n=305,742 controls) highlighted 27 and 8 loci, respectively, of which 13 were independent from the 78 sleep duration loci (**Figure 1b, Supplementary Tables 7,8, Supplementary Figures 1e,f, Appendix 1**). Only the *PAX8* signal was shared across all 3 traits, consistently indicating associations between the minor allele and longer sleep duration. For most long sleep loci, we could exclude equivalent effects on short sleep based on non-overlapping effects (**Supplementary Figure 4**, **Supplementary Table 8**). Sensitivity analyses accounting for factors potentially influencing sleep did not alter the results (**Supplementary Table 9,10)** Future studies will be necessary to test if these loci reflect partially distinct genetic effects on short or long sleep, or reflect differences in statistical power in these dichotomized analyses.

**Figure 1b.**
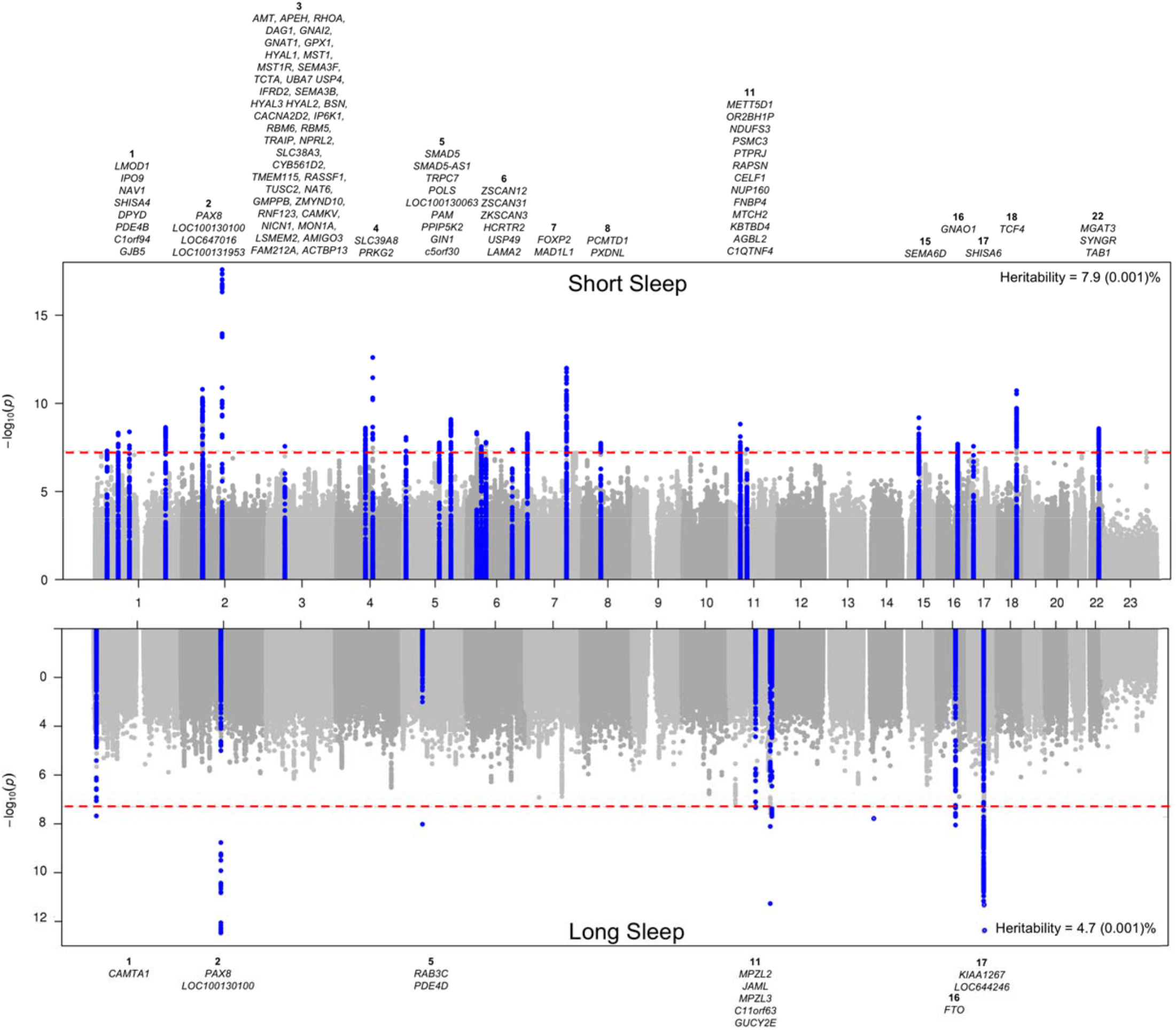
**B.** Short (cases *n* =106,192/305,742) and long (cases *n* =34,184/305,742) sleep.

We tested for independent replication of lead loci in the CHARGE consortium GWAS of adult sleep duration (*n* =47,180 from 18 studies^20^) and observed replication evidence for individual association signals at the *PAX8* and *VRK2* loci (*P* <6.4x10^-4^; **Supplementary Table 11,12, Supplementary Figure 5a**, **Appendix 1**). 55/70 signals showed a consistent direction of effect (binomial *P* =6.1x10^-7^), and a combined weighted genetic risk score (GRS) of the 70 signals was associated with a 0.66 [0.54-0.78] minutes longer sleep per allele (*P* =1.23x10^-25^) in the CHARGE study (**Table 1**). Consistently strong genetic correlation was observed between the CHARGE and UK Biobank studies (*r*_g_ (SE) =1.00 (0.123); *P*<0.001; **Supplementary Table 13**).

**Table 1.** A risk score of genetic variants for self-reported sleep duration (78 SNPs), self-reported short (27 SNPs) or long (8 SNPs) sleep duration associates with a) activity-monitor based measures of sleep fragmentation, timing and duration from 7 day accelerometry in the UK Biobank (n =85,499), b) self-reported sleep duration in the CHARGE study (n =47,180) and c) self-reported sleep duration in the EAGLE study (n =10,554).

| Study | Trait | Sleep Duration GRS |  | Short Sleep GRS |  | Long Sleep GRS |  |
| --- | --- | --- | --- | --- | --- | --- | --- |
|  |  | Beta or OR* [95% CI] per effect allele | P value | Beta or OR* [95% CI] per effect allele | P value | Beta or OR* [95% CI] per effect allele | P value |
| **CHARGE Study (n =47,180)** | Self-reported sleep duration (minutes)# | **0.66 [0.54 – 0.78]** | **1.23 x 10<sup>-25</sup>** |  |  |  |  |
| **EAGLE Study (n =10,554)** | Self-reported sleep duration (minutes)* | **0.16 [0.02 – 0.30]** | **2.80 x 10<sup>-2</sup>** |  |  |  |  |
| **UK Biobank 7-day Accelerometry (n =85,499)** | *Sleep Duration Estimates* |  |  |  |  |  |  |
|  | Sleep duration (minutes) | **0.47 [0.40 – 0.53]** | **1.93 x 10<sup>-44</sup>** | **-0.43 [-0.56 – -0.31]** | **1.21 x 10<sup>-11</sup>** | **2.12 [1.65 – 2.59]** | **1.08 x 10<sup>-18</sup>** |
|  | Short sleep duration (n=13,760 cases, 66,110 controls) | **0.98 [0.98 – 0.99]\*** | **4.00 x 10<sup>-19</sup>** | **1.02 [1.01–1.02]\*** | **4.91 x 10<sup>-6</sup>** | **0.94 [0.92 – 0.97]\*** | **1.10 x 10<sup>-5</sup>** |
|  | Long sleep duration (n=5,629 cases, 66,110 controls) | **1.01 [1.01 – 1.02]\*** | **3.78 x 10<sup>-9</sup>** | 0.99 [0.98 – 1.00]* | 0.11 | **1.10 [1.07 – 1.14]\*** | **1.29 x 10<sup>-8</sup>** |
|  | Daytime Inactivity Duration (minutes) | **0.08 [0.03 – 0.13]** | **2.74 x 10<sup>-3</sup>** | 0.01 [-0.09 – 0.11] | 0.89 | **0.65 [0.28 – 1.02]** | **6.49 x 10<sup>-4</sup>** |
|  | Sleep duration standard deviation (minutes) | -0.02 [-0.07 – 0.02] | 0.34 | 0.05 [-0.04 – 0.14] | 0.26 | -0.07 [-0.40 – 0.27] | 0.69 |
|  | *Sleep Fragmentation Estimates* |  |  |  |  |  |  |
|  | Sleep efficiency % | **0.05 [0.04 – 0.06]** | **8.38 x 10<sup>-23</sup>** | **-0.05 [-0.07 – -0.04]** | **4.79 x 10<sup>-9</sup>** | **0.15 [0.08 – 0.22]** | **1.56 x 10<sup>-5</sup>** |
|  | Number of sleep bouts (n) | **0.02 [0.01 – 0.02]** | **1.59 x 10<sup>-10</sup>** | **-0.01 [-0.02 – 0.00]** | **2.42 x 10<sup>-3</sup>** | 0.02 [-0.01 – 0.05] | 0.24 |
|  | *Sleep Timing Estimates* |  |  |  |  |  |  |
|  | Midpoint of 5-hour daily period of minimum activity (L5 timing) (minutes) | -0.05 [-0.13 – 0.03] | 0.23 | 0.07 [-0.09 – 0.22] | 0.41 | 0.39 [-0.20 – 0.97] | 0.20 |
|  | Midpoint of 10-hour daily period of maximum activity (M10 timing) (minutes) | 0.03 [-0.06 – 0.12] | 0.51 | -0.05 [-0.23 – 0.12] | 0.55 | 0.65 [-0.02 – 1.32] | 6.00 x 10<sup>-2</sup> |
|  | Sleep Midpoint (minutes) | -0.03 [-0.07 – 0.01] | 0.20 | 0.01 [-0.07 – 0.08] | 0.88 | 0.05 [-0.24 – 0.34] | 0.74 |

Genetic risk scores for sleep duration, short sleep and long sleep were tested using the weighted genetic risk score calculated by summing the products of the sleep trait risk allele count for all 78, 27, or 8 genome-wide significant SNPs multiplied by the scaled effect from the primary GWAS using the GTX package in R. Effect estimates (Beta or OR) are reported per each additional effect allele for sleep duration, short sleep, or long sleep. Abbreviations: CI=confidence interval, GRS=genetic risk score, OR=odds ratio.

In the childhood/adolescent GWAS for sleep duration from the EAGLE consortium^28^ (*n* =10,554), marginal evidence of association was observed for the adult sleep duration loci, with 45/77 signals demonstrating consistent directionality (binomial *P* =0.031; **Supplementary Table 11,12, Supplementary Figure 5b, Appendix 1**). For a combined 77 SNP GRS, we observed an effect of 0.16 [0.02-0.30] minutes longer sleep per allele (*P* =0.03; **Table 1**). No significant overall genetic correlation was observed with GWAS of adult sleep duration (*r*_g_ (SE) =0.098 (0.076), *P* =0.20 with UK Biobank; **Supplementary Table 13**), as reported previously^28^, however because of known changes in sleep patterns throughout the lifespan^29–31^, larger GWAS of sleep duration in children/adolescents are needed. Results of a meta-analysis of all three sleep duration GWAS are shown in **Supplementary Table 11,12** (also **Supplementary Figure 5c**, **Appendix 1**).

Given the limitations of self-reported sleep duration^32,33^, we tested the 78 lead variants for association with 8 accelerometer-derived sleep estimates in a subgroup who had completed up to 7 days of wrist-worn accelerometry (*n* =85,499; **Supplementary Table 14**)^34^. The lead *PAX8* genetic variant was associated with 2.68 (0.29) min longer sleep duration (compared to 2.44 (0.16) min by self-report), 0.21 (0.04) % greater sleep efficiency, and 0.94 (0.23) min greater daytime inactivity duration per minor A allele (*P* <0.00064; **Supplementary Table 15**). The 5% of participants carrying the most of the 78 sleep duration-increasing alleles were estimated to have 9.7 [7.5-11.8] min longer sleep duration compared to the 5% carrying the fewest. The 78 SNP GRS associated with longer accelerometer-derived sleep duration, longer duration of daytime inactivity, greater sleep efficiency, and larger number of sleep bouts, but not with day-to-day variability in sleep duration or estimates of sleep timing (**Table 1**). A GRS of 27 short sleep variants was associated with shorter accelerometer-derived sleep duration, lower sleep efficiency and fewer sleep bouts, whereas a GRS of 8 long sleep variants associated with longer accelerometer-derived sleep duration, higher sleep efficiency, and longer daytime inactivity (**Table 1, Supplementary Table 16**).

The sleep duration association signals encompass >200 candidate causal genes and a summary of reported gene-phenotype annotations is shown in **Supplementary Table 17**. Compelling candidates include genes in the dopaminergic (*DRD2*, *SLC6A3*), MAPK/ERK signaling (*ERBB4*, *VRK2*, *KSR2*), orexin receptor (*HCRTR2*) and GABA (*GABRR1*) signaling systems^4,35^. Further, studies of sleep regulation in animal models prioritize several candidates (*GABRR1*, *GNAO1*, *HCRTR2*, *NOVA1*, *PITX3*, *SLC6A3* and *DRD2*, *VAMP2* for sleep duration; *PDE4B*, *SEMA3F* for short sleep; *PDE4D* for long sleep). Circadian genes within associated loci include *PER3*, *BTRC* and the previously implicated *PER1* ^36^, which may act through glucocorticoid stress related pathways to influence sleep duration. Association signals at 4 loci directly overlapped with other GWAS signals (*r*^2^ >0.8 in 1KG CEU; from NHGRI), with the shorter sleep allele associated with higher BMI (*FTO*), increased risk of Crohn’s disease (*NFKB1, SLC39A8, BANK1* region), febrile seizures and generalized epilepsy (*SCN1A*), and cardiometabolic risk (*FADS1/2* gene cluster), and decreased risk of interstitial lung disease (*MAPT*/*KANSL*). Fine-mapping using credible set analysis in PICS^37^ highlighted 52 variants (**Supplementary Tables 18,19**). Partitioning of heritability by functional annotations identified excess heritability across genomic regions conserved in mammals, consistent with earlier findings^24^, and additionally in regions with active promoters and enhancer chromatin marks (**Supplementary Table 20**).

Gene-based tests identified 235, 54 and 20 genes associated with sleep duration, short and long sleep, respectively (P ≤2.29x10^-6^; **Supplementary Table 21**, **22**). Pathway analyses of these genes using MAGMA^38^ and Pascal^39^ indicated enrichment of pathways including striatum and subpallium development, mechanosensory response, dopamine binding, catecholamine production, and long-term depression (**Figure 2a,b, Supplementary Table 23,24**). In agreement the *FADS1/2* signal, we also observe an enrichment in genes related to unsaturated fatty acid metabolism, supporting experimental and observational evidence linking polyunsaturated fatty acids with sleep and related diseases, including neuropsychiatric disorders and depression^40–42^.

**Figure 2.**
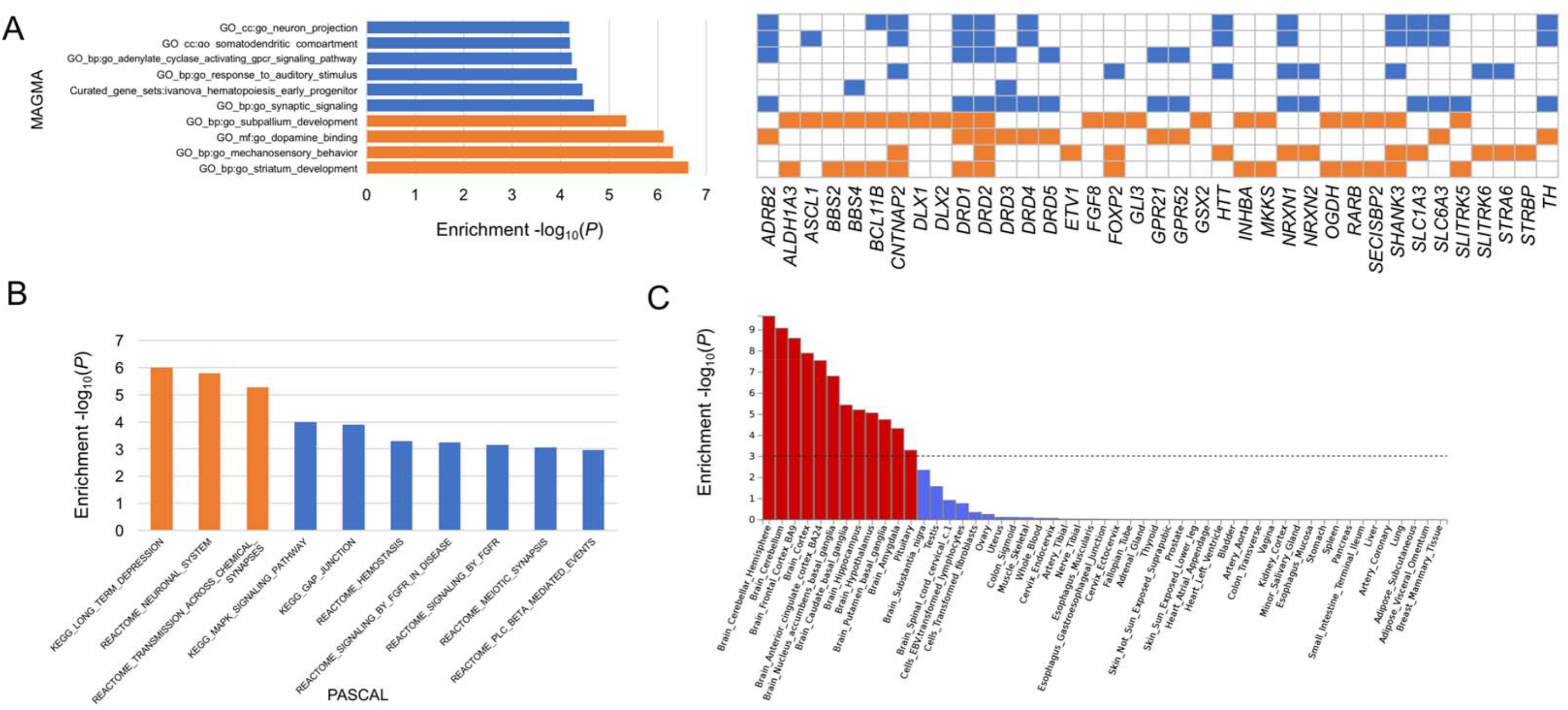
Pathway-based (A,B) and tissue expression enrichment analyses (C) for sleep duration. A) Pathway analysis is based on MAGMA gene-sets. Top 10 of 10,891 pathways are shown, and significant pathways are indicated in orange (*P* <4.59 x 10^-6^). For each significant pathway, respective sleep genes are indicated with a colored orange box. Sleep genes from significant pathways that overlap with remaining pathways are indicated in blue. B) Pathway analysis is based on Pascal (gene-set enrichment analysis using 1,077 pathways from KEGG, REACTOME, BIOCARTA databases) Top 10 pathways are shown, and significant pathways are indicated in orange (*P* <4.64 x 10^-5^). C) MAGMA tissue expression analysis using gene expression per tissue based on GTEx RNA-seq data for 53 specific tissue types. Significant tissues are shown in red (*P* <9.43 x 10^-4^). All pathway and tissue expression analyses in this figure can be found in tabular form in **Supplementary Tables 23,24,25**.

Tissue enrichment analyses of gene expression from GTEx tissues identified enrichment of associated genes in several brain regions including the cerebellum, a region of emerging importance in sleep/wake regulation^43^, frontal cortex, anterior cingulated cortex, nucleus accumbens, caudate nucleus, hippocampus, hypothalamus, putamen, and amygdala (**Figure 2c, Supplementary Table 25**). Enrichment was also observed in the pituitary gland. Integration of gene expression data with GWAS using transcriptome-wide association analyses in 11 tissues^44^ identified 38 genes for which sleep duration SNPs influence gene expression in the tissues of interest (**Supplementary Table 26**).

Several lead SNPs were associated with one or more of 3,144 human brain structure and function traits assessed in the UK Biobank (*P* <2.8x10^-7^, *n* =9,707; Oxford Brain Imaging Genetics Server^45^; **Supplementary Figure 6**). These include associations between the *PAX8* locus with resting-state fMRI networks (**Supplementary Figure 6a,h,m**), rs13109404 (*BANK1;* **Supplementary Figure 6b**) and bilateral putamen and striatum volume, possibly relating to functional findings on reward processing after experimental sleep deprivation^46^, and rs330088 (*PPP1R3B* region; **Supplementary Figure 6c**) and temporal cortex morphometry, which may relate to recent findings showing extreme sleep durations predict subsequent frontotemporal gray matter atrophy^47^.

Genome-wide genetic correlations using LD-score regression analyses^48^ indicated shared biological links between sleep duration and cognitive, psychiatric and physical disease traits (**Figure 3, Supplementary Table 27**). We observed positive genetic correlations between sleep duration and schizophrenia, bipolar disorder, and age at menarche, and a negative correlation with insomnia that persisted even upon excluding participants with psychiatric disorders, indicating that genetic relationships are not driven by the presence of co-morbid conditions. Both short and long sleep showed positive genetic correlations with depressive symptoms, waist circumference, waist-to-hip ratio and negative correlations with years of schooling. For short sleep, genetic correlations were also observed with insomnia, neuroticism, and smoking, and for long sleep, positive correlations were evident with schizophrenia, body fat, type 2 diabetes, and coronary artery disease.

**Figure 3.**
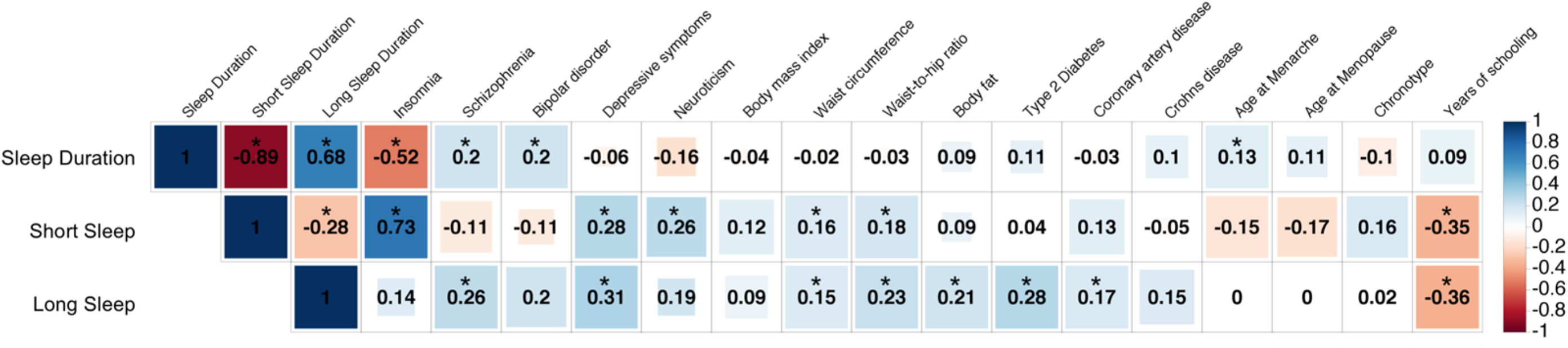
Genetic architecture shared between sleep duration and behavioral and disease traits. LD score regression estimates of genetic correlation (r_g_) were obtained by comparing genome-wide association estimates for sleep duration with summary statistics estimates from 224 publically available GWAS. Blue, positive genetic correlation; red, negative genetic correlation; rg values are displayed for significant correlations. Larger colored squares correspond to more significant *P* values, and asterisks indicate significant (*P* <2.2 x 10^-4^) genetic correlations. All genetic correlations in this report can be found in tabular form in **Supplementary Table 27**.

Mendelian Randomization (MR) analyses to test for causal links between sleep duration and genetically correlated traits suggested longer sleep duration is causal for increased risk of schizophrenia [weighted median: 0.008 (0.003) *P* =4.29 x 10^-3^; inverse variance weighted: 0.008 (0.003) *P* =1.82 x 10^-2^], consistent with previous findings^24,49,50^ (**Figure 4, Supplementary Table 28**). No other causal links were identified. Furthermore, no evidence of causal association was evident between sleep duration (excluding *FTO*) and BMI from the GIANT and UK Biobank datasets or with type 2 diabetes from the DIAGRAM and UK Biobank datasets (**Supplementary Table 28**, **Supplementary Figure 7**).

**Figure 4.**
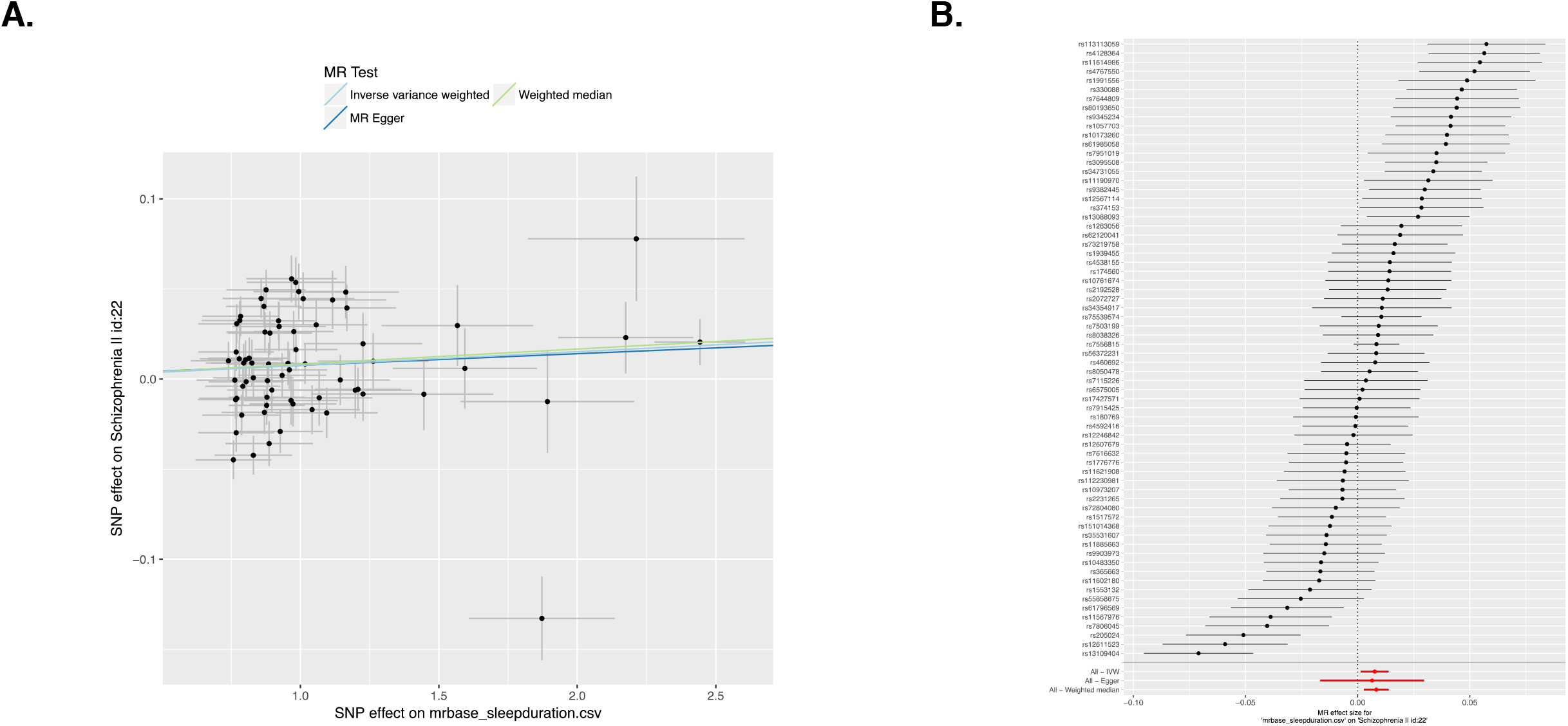
Causal relationship of sleep duration with schizophrenia in the UK Biobank. Association between single nucleotide polymorphisms (SNP) associated with sleep duration and schizophrenia (A) and forest plot shows the estimate of the effect of genetically increased sleep duration on schizophrenia (B). Results are shown for three MR association tests. Forest plots show each SNP with the 95% confidence interval (gray line segment) of the estimate and the Inverse Variance MR, MR-Egger, and Weighted Median MR results in red. MR results can be found in tabular form in **Supplementary Table 28**.

In summary, our GWAS for sleep duration expands our understanding of the genetic architecture of sleep duration, identifying 76 additional independent loci beyond the two previously known loci (*PAX8* and *VRK2*^20,23,24^). Further replication is important. We observed largely consistent effects with accelerometer-estimated sleep duration even though self-report, actigraphy and polysomnography estimated sleep duration provide both unique and overlapping information, have different sources of measurement error and may reflect different neurophysiological and psychological aspects^32,33^. Association of the sleep duration GRS with sleep efficiency suggests that sleep duration genetic loci might impact other correlated parameters such as sleep latency, sleep fragmentation and early awakening. Investigation of a role of these loci in EEG-derived physiological correlates of sleep architecture and sleep homeostasis from polysomnography, and follow-up in cellular and animal models will help to dissect functional mechanisms. All together, this work and follow-up studies will advance understanding of the molecular processes underlying sleep regulation and should open new avenues of treatment for sleep and related disorders.

## Online Methods

### Population and study design

Study participants were from the UK Biobank study, described in detail elsewhere^51^. In brief, the UK Biobank is a prospective study of >500,000 people living in the United Kingdom. All people in the National Health Service registry who were aged 40-69 and living <25 miles from a study center were invited to participate between 2006-2010. In total 503,325 participants were recruited from over 9.2 million invitations. Extensive phenotypic data were self-reported upon baseline assessment by participants using touchscreen tests and questionnaires and at nurse-led interviews. Anthropometric assessments were also conducted and health records were obtained from secondary care data from linked Hospital episode statistics (HES) obtained up until 04/2017. For the current analysis, 24,533 individuals of non-white ethnicity (as defined in “Genotyping and quality control”) were excluded to avoid confounding effects.

### Sleep duration and covariate measures

Study participants (*n* ~500,000) self-reported sleep duration at baseline assessment. Participants were asked, “About how many hours sleep do you get in every 24 hours? (please include naps),” with responses in hour increments. Sleep duration was treated as a continuous variable and also categorized as either short (6 hours or less), normal (7 or 8 hours), or long (9 hours or more) sleep duration. Extreme responses of less than 3 hours or more than 18 hours were excluded^23^ and “Do not know” or “Prefer not to answer” responses were set to missing. Participants who self-reported any sleep medication (see **Supplementary Method 1**) were excluded. Furthermore, participants who self-reported any shift work or night shift work or those with prevalent chronic disease (*i.e*., breast, prostate, bowel or lung cancer, heart disease or stroke) or psychiatric disorders (see **Supplementary Method 2**) were later additionally excluded in a secondary GWAS.

Participants further self-reported age, sex, caffeine intake (self-reported cups of tea per day and cups of coffee per day), daytime napping (“Do you have a nap during the day?”), smoking status, alcohol intake frequency (never, once/week, 2-3 times/week, 4-6 times/week, daily), menopause status, and employment status during assessment. Socio-economic status was represented by the Townsend deprivation index based on national census data immediately preceding participation in the UK Biobank. Weight and height were measured and body-mass index (BMI) was calculated as weight (kg) / height^2^(m^2^). Cases of sleep apnea were determined from self-report during nurse-led interviews or health records using International Classification of Diseases (ICD)-10 codes for sleep apnea (G47.3). Cases of insomnia were determined from self-report to the question, “Do you have trouble falling asleep at night or do you wake up in the middle of the night?" with responses “never/rarely”, “sometimes”, “usually”, “prefer not to answer”. Participants who responded “usually” were set as insomnia cases, and remaining participants were set as controls. Missing covariates were imputed by using sex-specific median values for continuous variables (*i.e*., BMI, caffeine intake, alcohol intake, and Townsend index), or using a missing indicator approach for categorical variables (*i.e*., napping, smoking, menopause, and employment).

### Activity-monitor derived measures of sleep

Actigraphy devices (Axivity AX3) were worn 2.8 - 9.7 years after study baseline by 103,711 individuals from the UK Biobank for up to 7 days. Details are described elsewhere^52^. Of these 103,711 individuals, we excluded 11,067 individuals based on accelerometer data quality. Samples were excluded if they satisfied at least one of the following conditions (see also http://biobank.ctsu.ox.ac.uk/crystal/label.cgi?id=1008): a non-zero or missing value in data field 90002 (“Data problem indicator”), “good wear time” flag (field 90015) set to 0 (No), “good calibration” flag (field 90016) set to 0 (No), “calibrated on own data” flag (field 90017) set to 0 (No) or overall wear duration (field 90051) less than 5 days. Additionally, samples with extreme values of mean sleep duration (<3 hours or >12 hours) or mean number of sleep periods (<5 or >30) were excluded. 85,502 samples remained after non-white ethnicity exclusions. Sleep measures were derived by processing raw accelerometer data (.cwa). First we converted .cwa files available from the UK Biobank to .wav files using Omconvert (https://github.com/digitalinteraction/openmovement/tree/master/Software/AX3/omconvert) for signal calibration to gravitational acceleration^52,53^ and interpolation^52^. The .wav files were processed with the R package GGIR to infer activity monitor wear time^54^, and extract the z-angle across 5-second epoch time-series data for subsequent use in estimating the sleep period time window (SPT-window)^34^ and sleep episodes within it^55^.

The SPT-window was estimated using an algorithm described in^34^, implemented in the GGIR R package and validated using PSG in an external cohort. Briefly, for each individual, median values of the absolute change in z-angle (representing the dorsalventral direction when the wrist is in the anatomical position) across 5-minute rolling windows were calculated across a 24-hour period, chosen to make the algorithm insensitive to activity-monitor orientation. The 10^th^ percentile was incorporated into the threshold distinguishing movement from non-movement. Bouts of inactivity lasting ≥30 minutes are recorded as inactivity bouts. Inactivity bouts that are <60 minutes apart are combined to form inactivity blocks. The start and end of longest block defines the start and end of the SPT-window^34^. *Sleep duration*. Sleep episodes within the SPT-window were defined as periods of at least 5 minutes with no change larger than 5° associated with the z-axis of the accelerometer, as motivated and described^55^. The summed duration of all sleep episodes was used as indicator of sleep duration. *Sleep efficiency*. This was calculated as sleep duration (defined above) divided by the time elapsed between the start of the first inactivity bout and the end of the last inactivity bout (which equals the SPT-window duration). *Number of sleep bouts within the SPT-window*. This is defined as the number of sleep bouts separated by last least 5 minutes of wakefulness within the SPT-window. The least-active five hours (*L5*) and the most-active ten hours (*M10*) of each day were defined using a five-hour and ten-hour daily period of minimum and maximum activity, respectively. These periods were estimated using a rolling average of the respectively time window. L5 was defined as the number of hours elapsed from the previous midnight whereas M10 was defined as the number of hours elapsed from the previous midday. *Sleep midpoint* was calculated for each sleep period as the midpoint between the start of the first detected sleep episode and the end of the last sleep episode used to define the overall SPT-window (above). This variable is represented as the number of hours from the previous midnight, e.g. 2am = 26. *Daytime inactivity duration* is the total daily duration of estimated bouts of inactivity that fall outside of the SPT-window. All activity-monitor phenotypes were adjusted for age at accelerometer wear, sex, season of wear, release (categorical; UK BiLeVe, UK Biobank Axiom interim, release UK Biobank Axiom full release) and number of valid recorded nights (or days for M10) when performing the association test in BOLT-LMM. Genetic risk scores for sleep duration, short sleep and long sleep were tested using the weighted genetic risk score calculated by summing the products of the sleep trait risk allele count for all 78, 27, or 8 genome-wide significant SNPs multiplied by the scaled effect from the primary GWAS using the GTX package in R^56^.

### Genotyping and quality control

Phenotype data is available for 502,631 subjects in the UK Biobank. Genotyping was performed by the UK Biobank, and genotyping, quality control, and imputation procedures are described in detail here^57^. In brief, blood, saliva, and urine was collected from participants, and DNA was extracted from the buffy coat samples. Participant DNA was genotyped on two arrays, UK BiLEVE and UK Biobank Axiom with >95% common content and genotypes for ~800,000 autosomal SNPs were imputed to two reference panels. Genotypes were called using Affymetrix Power Tools software. Sample and SNPs for quality control were selected from a set of 489,212 samples across 812,428 unique markers. Sample QC was conducted using 605,876 high quality autosomal markers. Samples were removed for high missingness or heterozygosity (968 samples) and sex chromosome abnormalities (652 samples). Genotypes for 488,377 samples passed sample QC (~99.9% of total samples). Marker based QC measures were tested in the European ancestry subset (n=463,844), which was identified based on principal components of ancestry. SNPs were tested for batch effects (197 SNPs/batch), plate effects (284 SNPs/batch), Hardy-Weinberg equilibrium (572 SNPs/batch), sex effects (45 SNPs/batch), array effects (5417 SNPs), and discordance across control replicates (622 on UK BiLEVE Axiom array and 632 UK Biobank Axiom array) (p-value <10^-12^ or <95% for all tests). For each batch (106 batches total) markers that failed at least one test were set to missing. Before imputation, 805,426 SNPs pass QC in at least one batch (>99% of the array content). Population structure was captured by principal component analysis on the samples using a subset of high quality (missingness <1.5%), high frequency SNPs (>2.5%) (~100,000 SNPs) and identified the sub-sample of white British descent. We further clustered subjects into four ancestry clusters using K-means clustering on the principal components, identifying 453,964 subjects of European ancestry. Imputation of autosomal SNPs was performed centrally by the UK Biobank to UK10K haplotype, 1000 Genomes Phase 3, and Haplotype Reference Consortium (HRC) with the current analysis using only those SNPs imputed to the HRC reference panel. Autosomal SNPs were pre-phased using SHAPEIT3^58^ and imputed using IMPUTE4. In total ~96 million SNPs were imputed. Related individuals were identified by estimating kinship coefficients for all pairs of samples, using only markers weakly informative of ancestral background. In total there are 107,162 related pairs comprised of 147,731 individuals related to at least one other participants in the UK Biobank.

### Genome-wide association analysis

Genetic association analysis was performed in related subjects of European ancestry (*n* =446,118) using BOLT-LMM^25^ linear mixed models and an additive genetic model adjusted for age, sex, 10 principal components of ancestry, genotyping array and genetic correlation matrix [jl2] with a maximum per SNP missingness of 10% and per sample missingness of 40%. We used a genome-wide significance threshold of 5x10^-8^ for each GWAS. Genetic association analysis was also performed in unrelated subjects of white British ancestry (n=326,224) using PLINK^59^ linear/logistic regression and an additive genetic model adjusted for age, sex, 10 PCs and genotyping array to determine SNP effects on sleep traits. We used a hard-call genotype threshold of 0.1, SNP imputation quality threshold of 0.80, and a MAF threshold of 0.001. Genetic association analysis for the X chromosome was performed using the genotyped markers on the X chromosome with the additional –sex flag in PLINK. Similarly, sex-specific GWAS were also performedusing BOLT-LMM^25^ linear mixed models. Trait heritability was calculated as the proportion of trait variance due to additive genetic factors measured in this study using BOLT-REML^25^, to leverage the power of raw genotype data together with low frequency variants (MAF≥0.001). Lambda inflation (λ) values were calculated using GenABEL in R^56^, and estimated values were consistent with those estimated for other highly polygenic complex traits. Additional independent risk loci were identified using the approximate conditional and joint association method implemented in GCTA (GCTACOJO)^60^.

### Post-GWAS analyses

#### Sensitivity analyses of top signals

Follow-up analyses on genome-wide significant loci in the primary analyses included covariate sensitivity analyses adjusting for BMI, insomnia, or caffeine intake adjustments individually, or a combined adjustment for BMI, day naps, Townsend index, smoking, alcohol intake, menopause status, employment status, and sleep apnea in addition to baseline adjustments for age, sex, 10 principal components of ancestry, and genotyping array. Sensitivity analyses were performed in PLINK 1.9^59^ using linear/logistic regression conducted only in unrelated subjects of white British ancestry.

#### Replication and meta-analyses with self-reported sleep duration GWAS

Using publicly available databases, we conducted a lookup of lead self-reported sleep duration signals in self-reported sleep duration GWAS results from adult (CHARGE) and childhood/adolescent (EAGLE). If lead signal was unavailable, a proxy SNP was used instead. In addition, we combined self-reported sleep duration GWAS results from adult (CHARGE) and childhood/adolescent (EAGLE) with the UK Biobank (primary model) in fixed-effects meta-analyses using the inverse variance-weighted method in METAL^61^. Meta-analyses were conducted first separately [UK Biobank + CHARGE (n=3,044,490 variants) or UK Biobank + EAGLE (n=7,147,509 variants)], then combined (UK Biobank + CHARGE + EAGLE; n=2,545,157 variants). A genetic risk score for sleep duration was tested using the weighted genetic risk score calculated by summing the products of the sleep duration risk allele count for as many available SNPs of the 78 genome-wide significant SNPs in each study (70 for CHARGE, 77 for EAGLE) multiplied by the scaled effect from the primary GWAS using the GTX package in R^56^.

#### Gene, pathway and tissue-enrichment analyses

Gene-based analysis was performed using Pascal^39^. Pascal gene-set enrichment analysis uses 1,077 pathways from KEGG, REACTOME, BIOCARTA databases, and a significance threshold was set after Bonferroni correction accounting for 1,077 pathways tested (*P* <0.05/1,077). Pathway analysis was also conducted using MAGMA^38^ geneset analysis in FUMA^62^, which uses the full distribution of SNP *P* values and is performed for curated gene sets and GO terms obtained from MsigDB (total of 10,891 pathways). A significance threshold was set after Bonferroni correction accounting for all pathways tested (*P* <0.05/10,891). Tissue enrichment analysis was conducted using FUMA^62^ for 53 tissue types, and a significance threshold was set following Bonferroni correction accounting for all tested tissues (*P* <0.05/53). Integrative transcriptome-wide association analyses with GWAS were performed using the FUSION TWAS package^44^ with weights generated from gene expression in 9 brain regions and 2 tissues from the GTEx consortium (v6). Tissues for TWAS testing were selected from the FUMA tissue enrichment analyses and here we present significant results that survive Bonferroni correction for the number of genes tested per tissue and for all 11 tissues.

#### Genetic correlation analyses

Post-GWAS genome-wide genetic correlation analysis of LD Score Regression (LDSC)^63–65^ using LDHub was conducted using all UK Biobank SNPs also found in HapMap3 and included publicly available data from 224 published genome-wide association studies, with a significance threshold after Bonferroni correction for all tests performed (*P* <0.05/224 tests). LDSC estimates genetic correlation between two traits from summary statistics (ranging from -1 to 1) using the fact that the GWAS effect-size estimate for each SNP incorporates effects of all SNPs in LD with that SNP, SNPs with high LD have higher X^2^ statistics than SNPs with low LD, and a similar relationship is observed when single study test statistics are replaced with the product of z-scores from two studies of traits with some correlation. Furthermore, genetic correlation is possible between case/control studies and quantitative traits, as well as within these trait types. We performed partitioning of heritability using the 8 pre-computed cell-type regions, and 25 pre-computed functional annotations available through LDSC, which were curated from large-scale robust datasets^63^. Enrichment both in the functional regions and in an expanded region (+500bp) around each functional class was calculated in order to prevent the estimates from being biased upward by enrichment in nearby regions. The multiple testing threshold for the partitioning of heritability was determined using the conservative Bonferroni correction (*P* <0.05/25 classes). Summary GWAS statistics will be made available at the UK Biobank web portal.

## Mendelian randomization analyses

MR analysis was carried out using MR-Base (https://www.biorxiv.org/content/early/2016/12/16/078972), using the inverse variance weighted approach as our main analysis method^66^, and MR-Egger^67^ and weighted median estimation^68^ as sensitivity analyses. MR results may be biased by horizontal pleiotropy – i.e. where the genetic variants that are robustly related to the exposure of interest (here sleep duration) independently influence levels of a causal risk factor for the outcome. IVW assumes that there is either no horizontal pleiotropy, or that, across all SNPs, horizontal pleiotropy is (i) uncorrelated with SNP-risk factor associations and (ii) has an average value of zero. MR-Egger assumes (i) but relaxes (ii) by explicitly estimating the non-zero mean pleiotropy, and adjusting the causal estimate accordingly. Estimation of the pleiotropy parameter means that the MR-Egger estimate is generally far less precise than the IVW estimate. The weighted median approach is valid if less than 50% of the weight is pleiotropic (i.e. no single SNP that contributes 50% of the weight or a number of SNPs that together contribute 50% should be invalid because of horizontal pleiotropy). Given these different assumptions, if all three methods are broadly consistent this strengthens our causal inference. For all our MR analyses we used two-sample MR, in which, for all 78 GWAS hits identified in this study for sleep duration, we looked for the per allele difference in odds (binary outcomes) or means (continuous) with outcomes from summary publicly available data in the MR-Base platform. Results are therefore a measure of ‘longer sleep duration’ and sample 1 is UK Biobank (our GWAS) and sample 2 a number of different GWAS consortia covering the outcomes we explored (**Supplementary Table 31,32**). The number of SNPs used in each MR analysis varies by outcome from because of some SNPs (or proxies for them) not being located in the outcome GWAS.

## Acknowledgements and Funding

This research has been conducted using the UK Biobank Resource. We would like to thank the participants and researchers from the UK Biobank who contributed or collected data. ARW is supported by 323195:SZ-245 50371-GLUCOSEGENES-FP7-IDEAS-ERC. BC is supported by K01-HL135405-01, R01-HL113338-04, R35-HL135818-01, American Thoracic Society Foundation Unrestricted Grant (Sleep). DG is supported by NIH/NHLBI contracts N01☐HC☐25195 and N02☐HL☐6☐4278 and cooperative agreement U01HL53941. DWR is supported by Wellcome Investigator award 107849/Z/15/Z. HT is supported by Dutch Medical Research Foundation grants (016.VICI.170.200 and VIDI 017.106.370). JML is supported by F32DK102323, 4T32HL007901. MG is supported by the Spanish Government of Investigation, Development and Innovation (SAF2017-84135-R) including FEDER co-funding, and NIDDK R01DK105072. MKR is supported by The University of Manchester Research Infrastructure Fund. MKR has acted as a consultant for GSK, Novo Nordisk, Roche and MSD, and also participated in advisory board meetings on their behalf. MKR has received lecture fees from MSD and grant support from Novo Nordisk, MSD and GSK. MNW is supported by Wellcome Trust Institutional Strategic Support Award (WT097835MF). SEJ is funded by the Medical Research Council (grant: MR/M005070/1). SP is funded by MH108908(SP), HL135818(SR). TMF is funded by 323195:SZ-245 50371-GLUCOSEGENES-FP7-IDEAS-ERC. JT is funded by Diabetes Research and Wellness Foundation Fellowship. RB is funded by Wellcome Trust and Royal Society grant: 104150/Z/14/Z. HSD and RS are supported by NIH R01DK107859, (RS), NIH R01 DK102696 (RS, FAJLS), MGH Research Scholar Fund (RS) and NIH R35 HL135818 (SR).

## Author Contributions

The study was designed by HSD, SEJ, JML, VVH, DAL, MKR, MNW, and RS. HSD, SEJ, ARW, JML, VVH, JR, YS, KP, RB, JB, SK, ML, AIL, KS, JT, DG, HT, DWR, SP, TMF, SL, DAL, MKR, MNW, and RS participated in acquisition, analysis and/or interpretation of data. HSD, JML, HW and RS wrote the manuscript and all co-authors reviewed and edited the manuscript, before approving its submission. RS is the guarantor of this work and, as such, had full access to all the data in the study and takes responsibility for the integrity of the data and the accuracy of the data analysis.

## Competing financial interests

FAJLS has received speaker fees from Bayer Healthcare, Sentara Healthcare, Philips, Kellogg Company, and Vanda Pharmaceuticals. MKR reports receiving research funding from Novo Nordisk, consultancy fees from Novo Nordisk and Roche Diabetes Care, and modest owning of shares in GlaxoSmithKline

## Data Availability

Summary GWAS statistics will be made publicly available at the UK Biobank website (http://biobank.ctsu.ox.ac.uk/) and through the International Sleep Genetic Epidemiology Consortium (ISGEC) webpage (sleepgenetics.org).

